# BacScan: An Unbiased and Genome-Wide Approach to Identify Bacterial Highly Immunogenic Proteins

**DOI:** 10.1101/2023.07.26.550668

**Authors:** Junhua Dong, Qian Zhang, Jinyue Yang, Yacan Zhao, Zhuangxia Miao, Siyang Pei, Huan Qin, Guoyuan Wen, Anding Zhang, Pan Tao

## Abstract

Bacterial pathogens are the second leading cause of death worldwide. However, the development of bacterial vaccines has been challenged by the presence of multiple serotypes and the lack of cross-protection between serotypes. Therefore, there is an urgent need to identify protective antigens conserved across serotypes in order to develop a broadly protective vaccine. Here, we have developed an unbiased and genome-wide technique, BacScan, which uses bacterial-specific serum to rapidly identify highly conserved immunogenic proteins by combining phage display, immunoprecipitation, and next-generation sequencing. As a proof of concept, we identified 19 highly immunogenic proteins from *Streptococcus suis* core proteins. Immunoreactivity analysis of mouse, pig, and human sera indicated that 2 proteins could be the potential targets for the development of serological diagnostics. Eight proteins provided 20%-100% protection against *S. suis* challenge in immunized animals, indicating the potential vaccine targets. BacScan can be applied to any bacterial pathogen and has the potential to accelerate the development of a broadly protective bacterial vaccine.

**Teaser:** A novel method to identify the highly conserved immunogenic bacterial proteins as targets for the development a broadly protective bacterial vaccine.

## Introduction

Bacterial pathogens are the second leading cause of death worldwide after ischemic heart disease, and an estimated 13.6% of global deaths in 2019 were caused by the 33 clinically significant bacterial pathogens (1). Antimicrobials are widely used to treat or prevent bacterial pathogens, but overuse and misuse of antibiotics lead to antimicrobial resistance, which was associated with an estimated 4.95 million deaths in 2019 and has become an emerging threat to public health (2). Vaccination, which is the most effective strategy to control infectious diseases, has gained more attention for the prevention of bacterial infections. Bacteria have a high genetic diversity of strains, and each bacterial species usually has multiple serotypes, and most bacterial vaccines are less effective against heterologous serotypes (3–5). Therefore, a universal bacterial vaccine that provides protection against divergent serotypes within bacterial species is urgently needed to reduce the transmission of bacterial pathogens and slow down the development of antimicrobial resistance.

An efficient approach to developing broadly protective vaccines is to use the antigens that are conserved across different serotypes as vaccine targets. However, bacteria encode a large number of proteins, and it is difficult to identify the antigenic proteins that can induce protective immune responses. Many genome-wide screening strategies have been developed for antigen discovery, and each has its advantages and disadvantages (6–8). For example, reverse vaccinology uses in silico analysis of the bacterial genome to identify typically hundreds of candidate protein antigens, which are heterologously expressed and individually evaluated for vaccine potential in an animal model. This approach was first applied to *Neisseria meningitidis*, and three out of 350 candidate proteins were identified for the development of a subunit vaccine that is now licensed in 30 countries (9, 10). However, it is costly and time-consuming to express hundreds of antigens and evaluate their protective efficacy in animals. Filamentous phage libraries that randomly display bacterial proteins or peptides have been screened with bacterium-positive sera to identify highly immunogenic antigens, which specifically bind to sera and are enriched after multiple rounds of panning (11). However, filamentous phages assemble in the periplasm of *E. coli*, and expressed antigens must be transported to the periplasm to be displayed on phage capsids (12, 13). In addition, the expression of certain exogenous proteins can limit the proliferation of the recombinant phages. As a result, these phages are lost after multiple rounds of amplification in *E. coli*. Therefore, this traditional phage display may have a selection bias. Two-dimensional electrophoresis of *in vitro*-cultured bacteria coupled with western blot using bacterium-specific sera has also been used to identify immunogenic antigens (14–16). However, not all immunogenic proteins are expressed when bacteria are cultured *in vitro*, such as some toxins and secreted proteins. In addition, the proteins transferred to the blot membrane are denatured proteins that cannot be bound by antibodies that recognize conformational epitopes.

Here, by focusing on *Streptococcus suis*, we aimed to develop a novel unbiased genome-wide approach to discover the highly immunogenic antigens conserved across different serotypes for vaccine and diagnostic development. *S. suis* is an emerging zoonotic pathogen that has caused three human outbreaks since the first case of human infection was reported in Denmark in 1968 (17). Although most human infections have occurred in Asia, particularly in China and Southeast Asian countries, sporadic cases have been reported in 34 countries around the world (18). Swine are considered to be a natural reservoir of *S. suis*. Although some serotypes are pathogenic to pigs and cause meningitis, arthritis, and septicemia, *S. suis* is considered a commensal of pigs and commonly colonizes the upper respiratory tract of almost all pigs (19, 20). In fact, almost all reported human cases of *S. suis* infection have had close contact with pigs or raw pork (21–24). Recently, a large-scale phylogenomic analysis of *S. suis* isolates over 36 years showed that almost all human isolates clustered into a human-associated clade (HAC), which was further subdivided into three lineages, I, II, and III (25). Although it is not clear whether human-to-human transmission occurs through direct contact with patient materials, these studies suggest that the risk of human *S. suis* outbreaks is increasing. Antibiotics such as ceftriaxone and penicillin are used to treat *S. suis* infections in humans and swine (21). However, numerous antibiotic-resistance genes have been found in HAC lineage III strains (25), highlighting the urgent need for alternative approaches to prevent and treat *S. suis* infections.

Inactivated *S. suis* vaccines have been licensed for use in animals; however, the protection efficacy is serotype or strain dependent (26). In addition, no efforts have been reported to develop *S. suis* vaccines for human use. Therefore, a universal vaccine that provides protection against divergent serotypes is urgently needed to reduce *S. suis* transmission and slow down the development of antibiotic resistance. The identification of immunogenic antigens that are conserved across different serotypes is the first step in the development of broadly effective subunit vaccines to prevent *S. suis* infection in both pigs and humans. Many immunogenic proteins of *S. suis* have been identified, but many of these are not core genes and are absent in some serotypes or strains (26–30). Therefore, genome-wide identification of all immunogenic proteins and definition of vaccine antigens will greatly accelerate the development of universal *S. suis* vaccines for both animal and human use. Meanwhile, the identification of diagnostic targets from the immunogenic proteins also helps develop serological diagnostic techniques for the epidemiological surveys of *S. suis* in pigs. In particular, the HAC-specific diagnostic targets can identify pig populations infected with *S. suis* strains with zoonotic potential for human infection, which will greatly reduce the risk of *S. suis* outbreaks in humans.

Here, we developed an unbiased and genome-wide technique, BacScan, to rapidly identify all the immunogenic core proteins of *S. suis* by combining phage display library, phage-immunoprecipitation, phage-PCR, and next-generation sequencing. BacScan uses T7 instead of filamentous phage and single round of selection instead of multi-round panning to avoid the biased selection of traditional phage display methods. Phage-PCR is used to amplify all the enriched phages, and the resulting PCR products are analyzed by next-generation sequencing and data mining to identify all the highly immunogenic proteins (HIPs) in a single round of selection. As a proof of concept, we identified 19 HIPs from all 793 core proteins of *S. suis*. Immunoreactivity analysis of mouse, pig, and human sera indicated that 2 of these HIPs could be the potential targets for the development of serological diagnostics. When administered intraperitoneally, 8 HIPs were able to provide animals with 20%-100% protection against *S. suis* challenge. Our studies have demonstrated that BacScan is a powerful approach to rapidly identify immunogenic proteins conserved across bacterial serotypes for the development of vaccines and diagnostics, which are highly desired to reduce the transmission of bacterial pathogens and slow the development of antimicrobial resistance.

## Results

### A BacScan platform for the unbiased and genome-wide screening of *S. suis* highly immunogenic proteins (HIPs)

Inspired by VirScan (31), a detailed experimental scheme was developed to identify bacterial immunogenic proteins in a genome-wide and unbiased manner, which we call BacScan (Fig. 1A). To identify antigenic proteins that present in all *S. suis* strains, the core genome containing 793 CDSs was identified and used together with 12 genes encoding previously identified antigenic proteins to construct a T7 phage library. All the 805 CDSs were split into 2,060 fragments of 600 bp with 300 bp overlap. The fragments were amplified from *S. suis* SC19 genome by PCR and inserted into T7 genome to generate a phage display library, which was used to enrich for phages recognized by *S. suis*-specific sera. The antibody-bound phages were captured by protein A/G coated beads, and unbound phages were removed by multiple washes. The *S. suis* gene fragments in the genome of enriched T7 phages were amplified by PCR using a pair of phage-specific primers, and the generated PCR products were sequenced by next-generation sequencing (NGS). The 600-bp fragments were used instead of full-length genes because homogenous-length fragments can significantly reduce the bias of PCR amplification. In addition, since the average size of the protein structure domain is about 200 amino acids (32), we hypothesized that 600-bp fragments could preserve protein structure intact to the greatest extent. After NGS sequencing, all the HIPs can be identified by analyzing the read count for each fragment before and after enrichment using the model described in VirScan (31).

**Figure 1.**
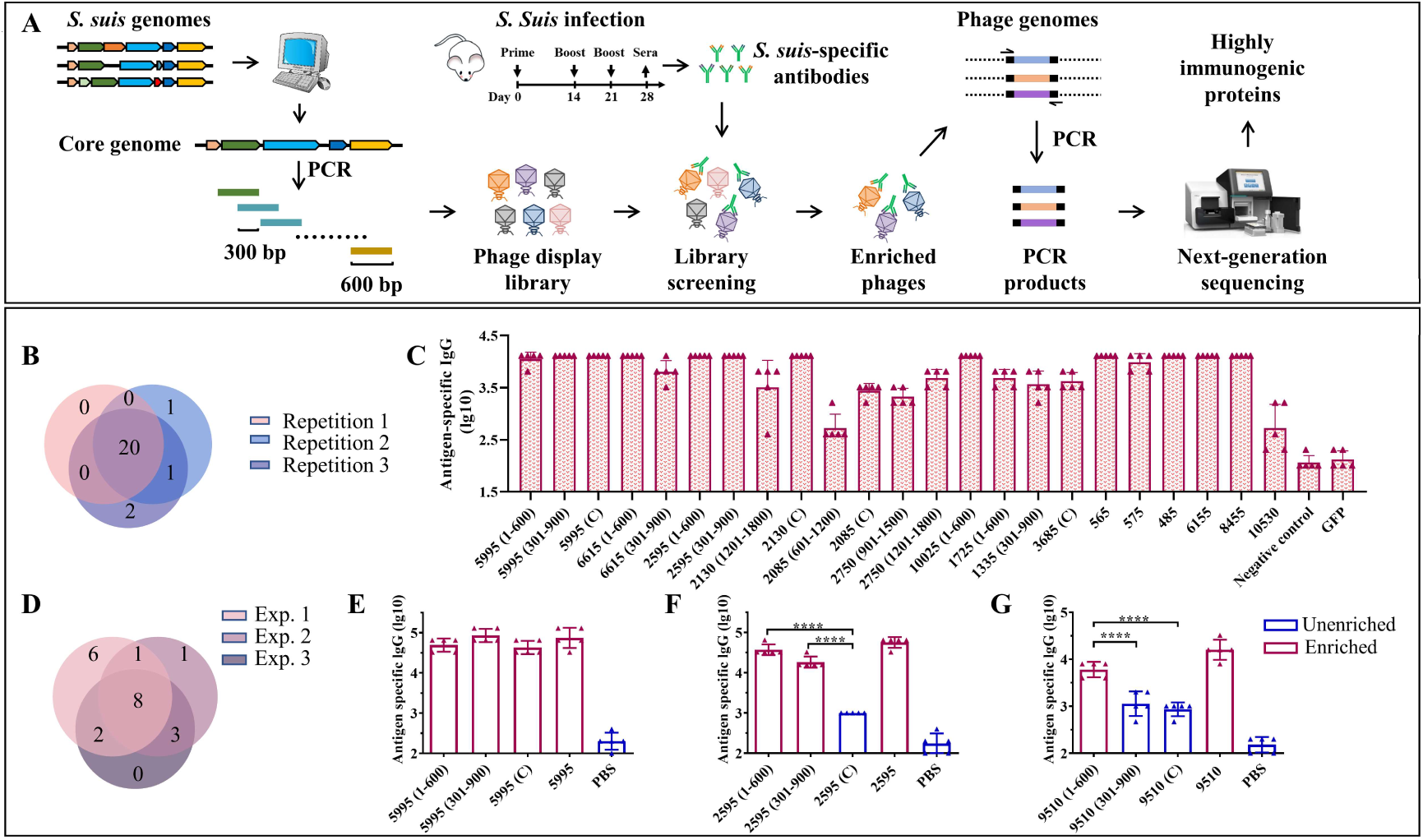
A BacScan platform for screening HIPs of *S. suis*. **(A)** Schematic diagram of BacScan for screening of HIPs from *S. suis* core genes. The *S. suis* core genes were identified using bioinformatic analysis of *S. suis* genomes and split into fragments of 600 bp in length, with 300 bp overlap between adjacent fragments by PCR. The fragments were inserted into a T7 vector to generate a T7 phage display library, which was incubated with *S. suis* specific sera. Phages bound by *S. suis*-specific antibodies were captured using protein A/G beads. The *S. suis* gene fragments in the T7 phage genome were amplified and sequenced by next-generation sequencing. The HIPs were identified by the phip-stat program. **(B)** The Venn diagram shows the screening results of three parallel experiments. A total of 24 fragments were identified, and 20 of them were enriched in three parallel experiments. The number in brackets indicates the location of the fragment in the gene. **(C)** ELISA results show that 23 enriched fragments were recognized by *S. suis*-specific sera. The GFP protein and *S. suis*-negative mouse sera were used as controls. **(D)** The Venn diagram shows the screening results of three independent experiments using the same serum sample. **(E-G)** Identification of the highly immunogenic regions of proteins 5995 **(E)**, 2595 **(F)**, and 9510 **(G)** by BacScan.

To prepare *S. suis*-specific sera, mice were infected three times by intraperitoneal (i.p.) injection of 4×10^7^ CFU *S. suis* SC19 strain, and sera were collected to confirm the presence of *S. suis*-specific IgG by ELISA (Fig. 1A and Fig. S1A). Approximately 2 μg of IgG per sample was used to enrich phages displaying *S. suis* immunogenic proteins. Each sample was run in triplicate to assess the reproducibility of the BacScan. A total of 24 gene fragments belonging to 17 genes were identified, 20 of which were enriched in three parallel experiments (Fig. 1B), demonstrating the reproducibility of the BacScan. Among these potential HIPs, 2750, 5995, 10025, and 6615, have been reported in previous studies (33–36). We attempted to express all of these gene fragments in *E. coli* and were able to purify 23 recombinant proteins. Since the 10530 fragment (301-900) tends to aggregate during purification, we expressed the full-length 10530 protein instead of the fragment. The binding activities of these proteins to *S. suis*-specific sera were determined using ELISA (Fig. 1C and Fig S1B). The GFP protein and negative mouse sera were used as controls. All the 23 recombinant proteins but not the GFP control can be recognized by *S. suis*-specific sera (Fig. 1C). However, the IgG endpoint titers of different proteins/fragments are variable. Titers of anti-2085 (601-1200) and anti-10530 IgG are lower than other HIPs specific IgG. (Fig. 1C). Four fragments 1335 (301-900), 2750 (901-1500), 6615 (301-900), and 1725 (1-600), which were enriched in only one or two of the three parallel experiments, can also be recognized by *S. suis*-specific sera. To avoid missing HIPs, we considered the fragments as long as they were enriched in at least one of the three parallel experiments.

To determine inter-experimental variation, BacScan was performed on the same serum sample in two additional independent experiments (Fig. S2). 13 HIPs were identified in both experiments 2 and 3, 11 of which were shared (Fig. 1D). Of the total 21 HIPs identified (Table S1), 9 were shared between experiments 1 and 2, 10 were shared between experiments 1 and 3, and 8 were shared in all three experiments (Fig. 1D). In addition, each of the independent experiments was performed in triplicate, and most of the HIPs were identified in three parallel replicates (Fig. S2A to S2C), further demonstrating the reproducibility of the BacScan. These results suggest that there is variation in the BacScan between independent experiments. However, the number of HIPs identified for the same serum sample become increasingly saturated as the number of screens increased, and few new antigens were identified after three independent experiments (Fig. S2E).

Interestingly, we found that BacScan can identify not only immunogenic proteins, but also the highly immunogenic part of the protein (Fig. 1E to 1G). For example, immunogenic proteins 5995, 2595, and 9510 were all split into three fragments in the T7 library, and three out of nine fragments were not enriched in our BacScan screen (Fig. 1E to 1G). We expressed all the 9 fragments as well as the full-length proteins to compare their binding activities to *S. suis*-specific sera using ELISA (Fig. 1E-1G, Fig.S1B, and Fig. S2F). We found that the endpoint IgG titers of the enriched fragments were significantly higher than those of the unenriched fragments (Fig. 1E to 1G), indicating that BacScan can identify antigens immunodominant regions of antigens. To reduce the workload, we will use the full-length gene if a fragment of it is enriched in our selection, unless otherwise indicated.

### Characteristics of *S. suis* highly immunogenic proteins

To determine the characteristics of *S. suis* highly immunogenic proteins, we first performed COGs (Clusters of Orthologous Groups) analysis of the 805 proteins included in our phage library and obtained 20 annotated COGs (Fig. 2A, before screen). The 21 HIPs identified in the current study were grouped into 13 annotated COGs (Fig. 2A, after screen). In particular, the cluster M, which is related to the cell wall/membrane/envelope biogenesis, was significantly overrepresented (p=0.0037, Fisher Exact Test). The proportion of cluster M increased from 5.3% (before screen) to 23.8% (after screen) (Fig. 2B, pink dot). This could be explained by the fact that the surface-exposed proteins are more likely to be recognized by the immune system and therefore have a higher chance of being captured by *S. suis*-specific sera. In fact, the surface exposure is a key criterion of reverse vaccinology to predict vaccine antigens (37–41), and surface-exposed proteins were significant enriched in known bacterial vaccine antigens (42). The COG cluster S, whose function is unknown, was negatively associated with the HIPs (Fig. 2A and B).

**Figure 2.**
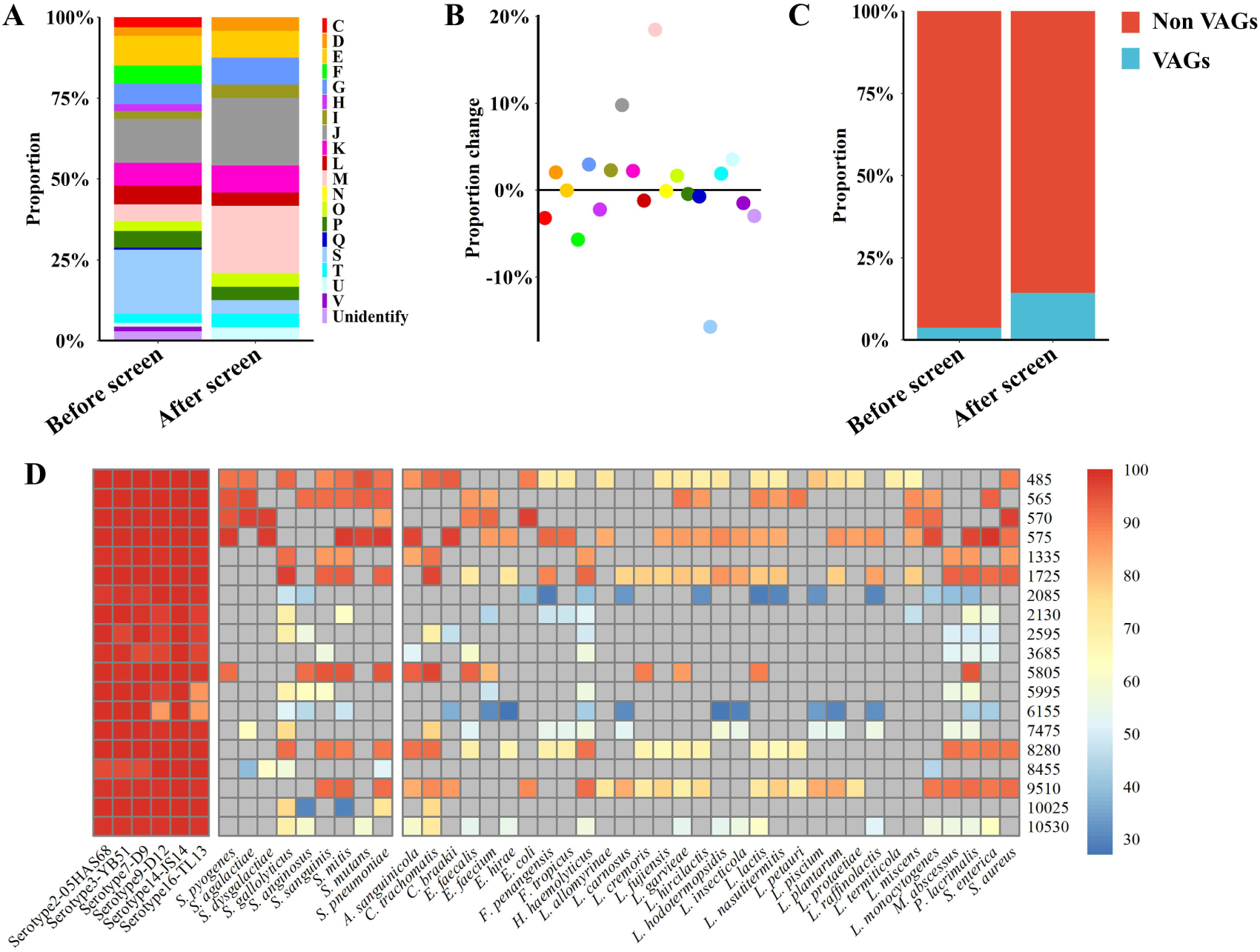
Characteristics of *S. suis* HIPs and their homology analysis. **(A)** and **(B)** Proteins related to the cell wall/membrane/envelope biogenesis were significantly enriched (p=0.0037, Fisher Exact Test). **(A)** The 805 proteins contained in the phage library were assigned to 20 annotated COGs, and the proportion of each COG before and after screening was shown. **(B)** The proportional changes of each COG before and after screening. **(C)** The proportion of VAGs is significantly increased after BacScan screening (p=0.0466, Fisher Exact Test). **(D)** Heatmap showing the conservation of 19 *S. suis* HIPs among different serotypes, representative human *streptococci* (see more in Supplementary Materials) and non-*streptococci* species. The color scale represents the conservation of each protein. All gray cells represent less than 80% coverage or no homologous proteins.

Antibodies directed against virulence factors can neutralize bacterial virulence and provide protection against infection. Therefore, virulence factors may be good vaccine targets. Previous studies have identified 71 virulence-associated genes (VAGs) from *S. suis* pan-genome (43), and 32 of them were included in our phage library. Interestingly, we found that the proportion of *S. suis* virulence-associated proteins were significantly enriched in the HIPs (p= 0.0466, Fisher Exact Test), increasing from 4.0% to 14.3 % (Fig. 2C).

The conserved antigens across different serotypes could be good targets for the development of broadly protective bacterial vaccines. Of the total 21 HIPs identified, two previously reported HIPs, 6615 and 2750, are not encoded by core genes and were therefore excluded from the subsequent sequence analysis. We found that all 19 HIPs encoded by core genes are highly conserved among different *S. suis* serotypes with 84-100% sequence identity (Fig. 2D), suggesting that they could be targets for the development of broadly protective *S. suis* vaccines. 15 HIPs also have 80%-98% sequence identity with proteins from other bacteria within or outside the genus *Streptococcus* (Fig. 2D and Fig. S3). Interestingly, two HIPs, 2085 and 6155, are highly conserved across serotypes but have relatively low homology with proteins from other bacteria (with less than 54% sequence identity) (Fig. 2D), suggesting that they may be specific targets for the development of serological techniques for the diagnosis of *S. suis*.

### Dynamic analysis of antibody repertoires to HIPs revealed diagnostic targets for serological detection of *S. suis*

To define which HIPs could be targets for development of vaccine and serologic diagnostics, we first determined the kinetics of antibody responses to the HIPs after *S. suis* infection. Four- to eight-week-old mice were infected once by intranasal (i.n.) administration with 8.0×10^7^ CFU *S. suis* SC19 strain, and sera were collected at different time points to determine HIP-specific IgG using ELISA. We attempted to express all the 21 HIPs in full-length using *E. coli* BL21, but failed to obtain the 1725, 570, 7475, and 2750 due to either solubility issues or weak binding to the HisTrap column (Fig. S1B, S2F and S4). The resulting 17 recombinant proteins were used individually as ELISA coating antigens. Most of the HIPs were able to induce antigen-specific IgG approximately 5 days after infection and persisted for at least 130 days (Fig. 3A). Overall, the reactivities of sera to HIPs become stronger with age. The 4-week-old mice were less efficient in producing HIP-specific IgG antibodies than 6-week-old and 8-week-old mice, which may be due to their immature immune systems. Although we were able to detect antibodies against all 17 HIPs, there was significant variation among the HIPs. We found that IgG antibodies were mainly directed against 13 HIPs, especially proteins 5995 and 10025 (Fig. 3A). The 5995-specific IgG was detected at 5 days, increased at 10 days, and peaked at 21 days post infection. Similarly, the 10025-specific IgG was detected at 5 days, increased at 7 days, and peaked at 10 days post infection (Fig. 3A). To determine whether the dose of infection affected the kinetics of antibody responses, 8-week-old mice were infected with 2.7×10^7^ (low), 8.0×10^7^ (medium), and 2.4×10^8^ (high) CFU *S. suis* SC19 strain, respectively. The dynamics of IgG antibody responses to the HIPs were similar between the medium- and high-dose groups. However, the development of HIP-specific IgG antibodies was delayed by 2-3 days in the low-dose group. Again, IgG antibodies were mainly directed against 13 HIPs, especially proteins 5995 and 10025 regardless of the infection dose (Fig. 3B).

**Figure 3.**
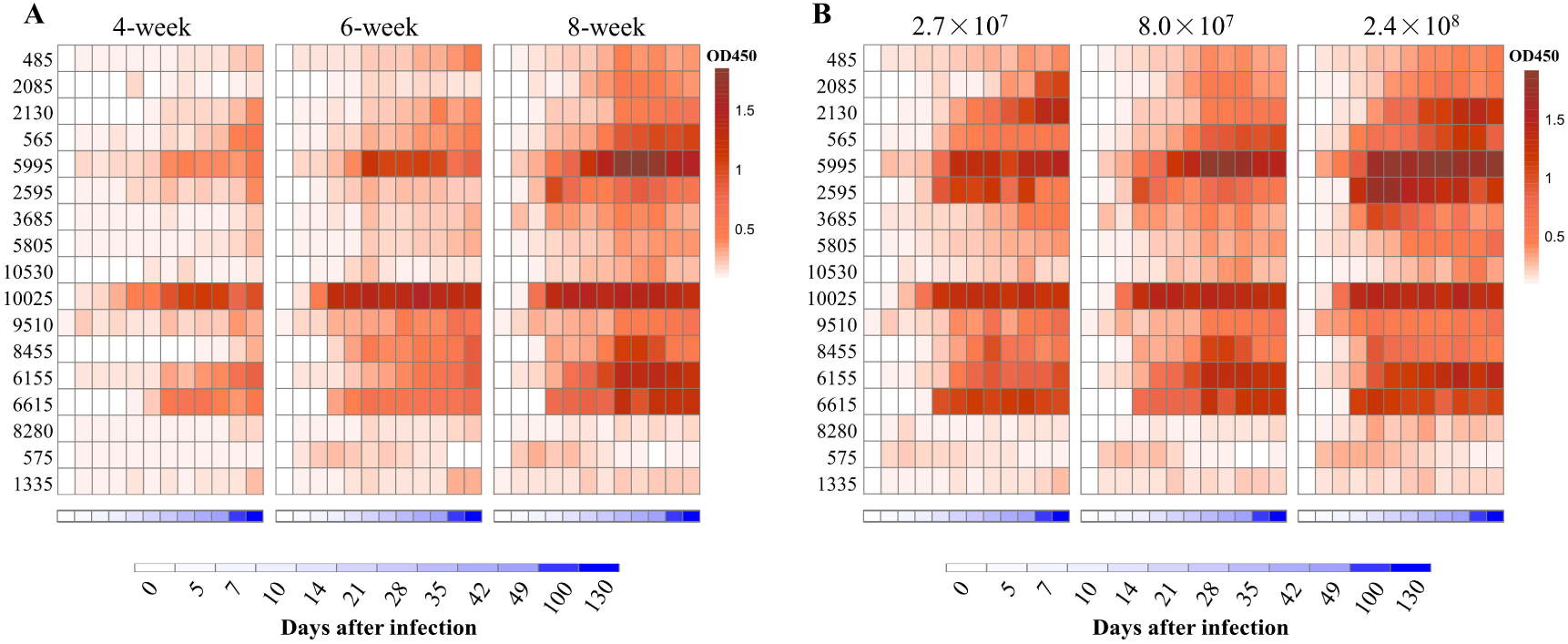
Dynamic analysis of antibody repertoires against HIPs after single infection with *S. suis* strain SC19 in mice. **(A)** Dynamic analysis of antibody repertoires in mice of different ages. Four- to eight-week-old mice were infected intranasally with 8.0×10^7^ CFU *S. suis* strain SC19. **(B)** Dynamic analysis of antibody repertoires in mice infected with different doses of *S. suis* SC19 strain. Eight-week-old mice were infected intranasally with 2.7×10^7^ (low dose), 8.0×10^7^ (medium dose), and 2.4×10^8^ (high dose) CFU *S. suis* SC19 strain, respectively. Sera were collected at different time points to determine HIP-specific IgG titers. Each row represents one HIP, and each column indicates the time after infection. The red color scale represents the O.D_450_ value of the ELISA. The blue color scale represents the time points.

We then focused on the top 10 HIPs that induced high IgG titers. To further determine the reactivities of pig sera to these HIPs, 20 clinical pig sera and 20 *S. suis*-negative sera were collected for ELISA analysis. As shown in Fig 4A, all the 10 HIPs can be recognized by clinical pig sera, although they exhibit variable binding activities. These results indicated that these pigs were likely infected with *S. suis* and further confirmed that these HIPs are highly immunogenic in pigs. However, since 6 of these HIPs showed high sequence identity (80-98%) to the proteins in other bacteria such as *E. coli*, *Salmonella enterica*, and *Staphylococcus aureus* (Fig. 2D and S3), we cannot exclude the possibility that these pigs were infected by other bacteria and that the ELISA signals were due to the cross-reactivity of the sera. As expected, *S. suis* negative sera did not react with these HIPs (Fig. 4A). Significantly, HIPs 2085 and 6155 which have low homology to proteins from other bacteria, showed high reactivities with clinical sera. In particular, the lowest value of O.D_450_ value in the clinical group is 4 times higher than that of the negative control for HIP 2085, making it an attractive target for the development of serological diagnostic methods.

**Figure 4.**
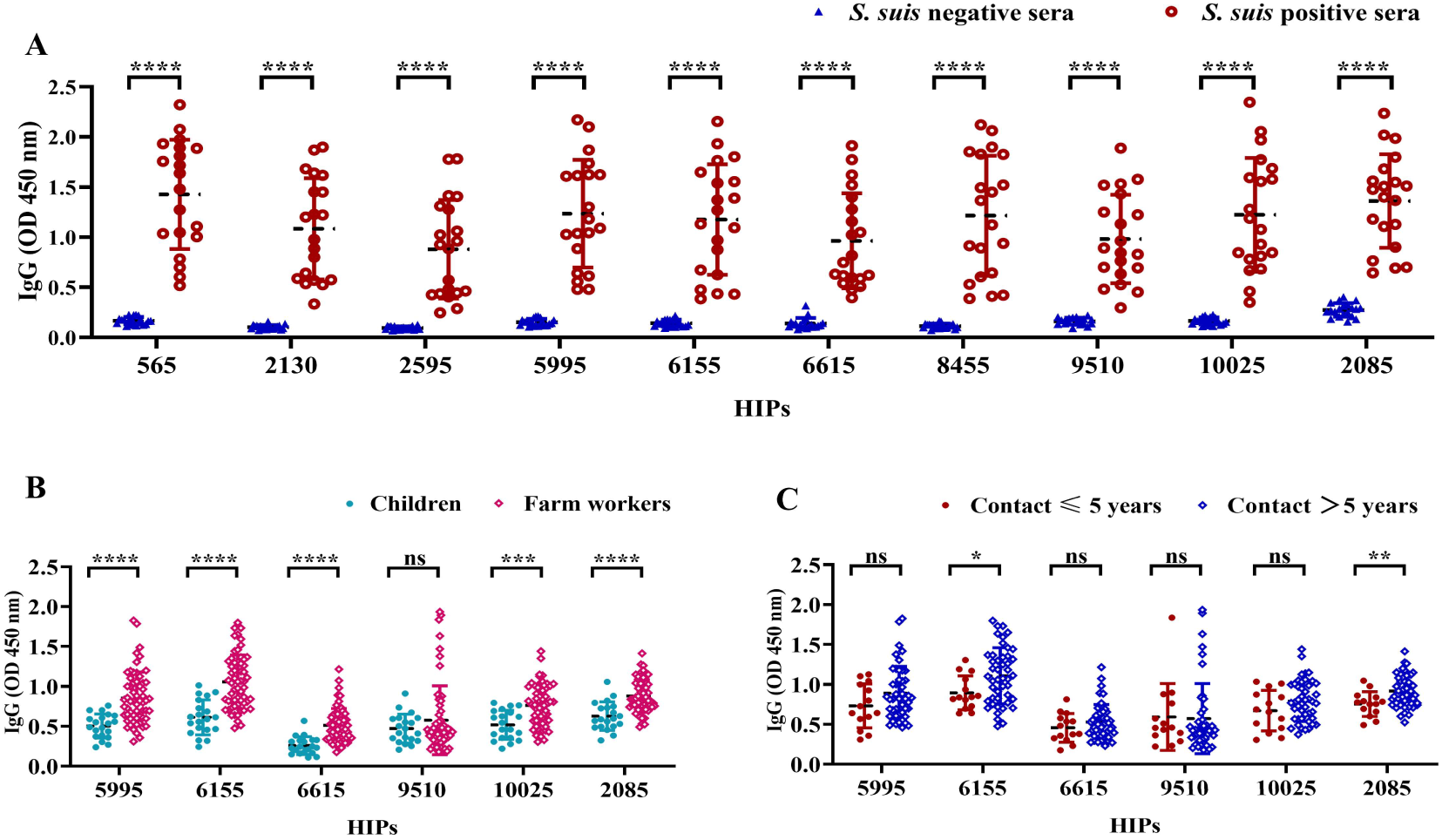
The reactivities of selected HIPs to swine and human sera. **(A)** The binding activities of HIPs to pig sera. Twenty clinical pig sera and 20 *S. suis*-negative sera were used to determine HIP-specific IgG by ELISA. Data were represented as mean ± S.D. ****P < 0.0001 (Student’s t-test). **(B)** ELISA results showing the binding activities of six HIPs to human sera. Sera from farm workers with a history of close contact with pigs (n=60) showed higher O.D_450_ than sera from children with no history of contact with pigs (n=20) for all tested HIPs except 9510. **(C)** Binding activities of six HIPs to sera from humans with varying lengths of history of exposure to pigs. HIPs 6155 and 2085 have higher binding activities to sera from humans with a history of > 5 years of close contact with pigs (n=46) compared to sera from humans with ≤ 5 years of history of exposure to pigs (n=14). Data were presented as mean ± S.D. *P < 0.05; **P < 0.01; ***P < 0.001; ****P < 0.0001 (Student’s t-test).

To determine whether HIPs 2085 and 6155 induce IgG antibodies in humans during *S. suis* infection, sera were collected for ELISA analysis from 60 farm workers with a history of close contact with pigs for 1-20 years. Sera from children with no history of contact with pigs were used as controls. HIPs 5995, 6615, 9510, and 10025 were also included in the serological analysis. The ELISA results showed that the sera from farm workers gave significantly higher O.D_450_ than the control sera for all tested HIPs except 9510 (Fig. 4B). Interestingly, the 2085- and 6155-specific O.D_450_ are significantly different between workers with more than 5 years and less than 5 years of experience in pig farming (Fig. 4C). These results suggest that HIPs 2085 and 6155 may be good targets for *S. suis* serological diagnostics in humans. However, more experiments are needed to further evaluate their potentialities.

### Identification of conserved protective antigens for the development of broadly protective subunit vaccines against *S. suis*

Three of the 21 HIPs identified, namely 2750 (known as Ide*_Ssuis_*), 6615 (known as SAO), and 5995 (known as PrsA), have been shown to induce protective immune responses in previous studies (33–35). Therefore, we randomly selected 5995 as a positive control and excluded 2750 and 6615 in our animal experiment. Due to the low yield of recombinant protein expressed in *E. coli*, HIPs 1725, 570, and 7475 were not included in our animal experiments. To define new HIPs capable of inducing protective immune responses for vaccine development, all the 16 recombinant HIPs were individually formulated with ISA 201 adjuvant and administered intraperitoneally to the mice twice a week (Fig. 5A). Sera were collected on days 0 and 28 for IgG titration, and mice were challenged with 6×10^9^ CFU *S. suis* SC19 strain 14 days after the boost. As shown in Fig. 5B, all 16 HIPs can induce antigen-specific IgG antibodies, whereas lower titers were observed for 1335-specific IgG. Mouse sera from the PBS control group did not react with any of these HIPs (ELISA plate was coated with a mixture of all 16 HIPs).

**Figure 5.**
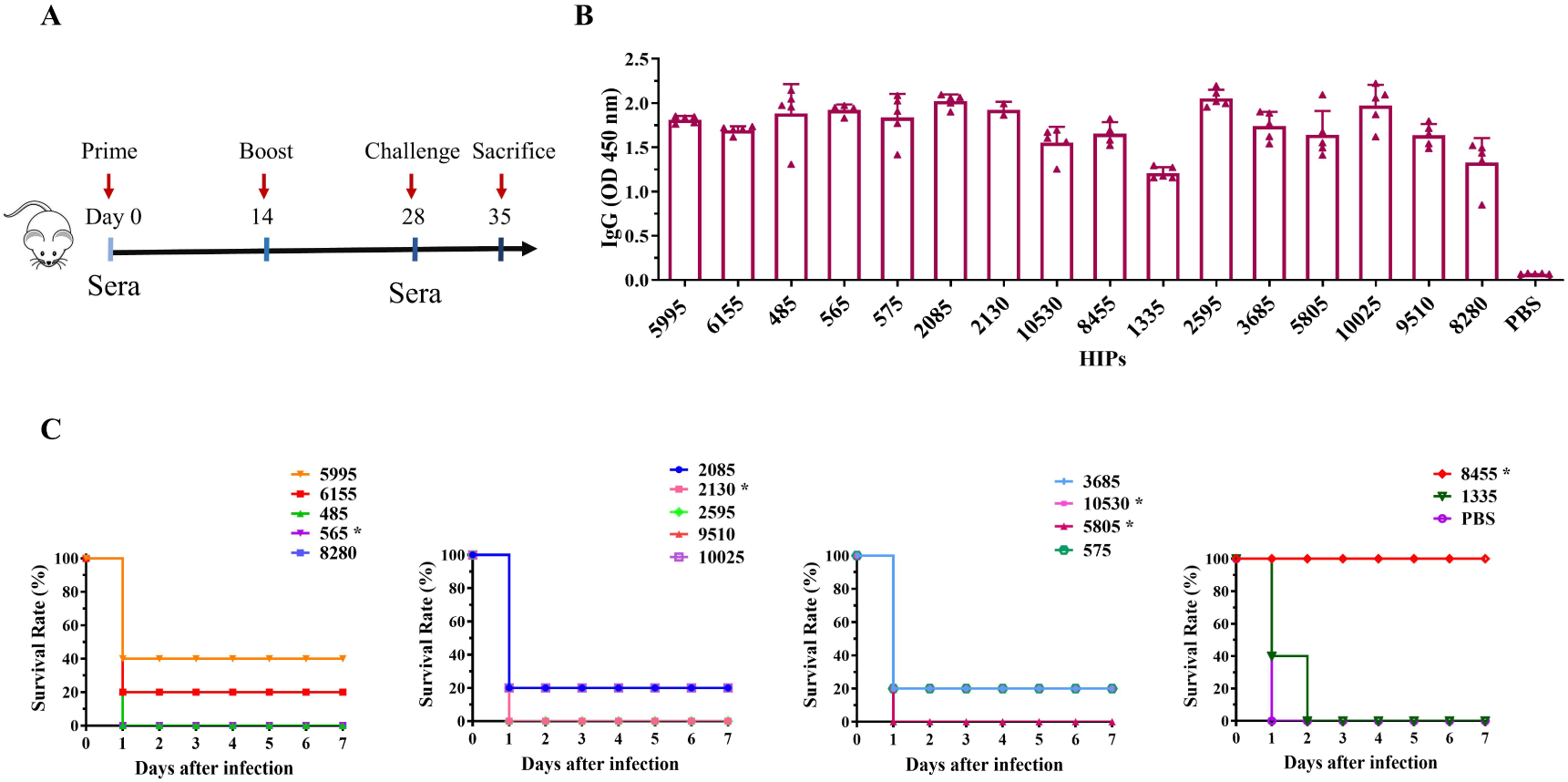
Immunogenicity and protective efficacy of HIPs in mice. **(A)** Mice were immunized twice by intraperitoneal injection, and sera were collected according to the scheme. **(B)** HIPs-specific IgG antibody titers were determined by ELISA. **(C)** The immune protection of each HIP was determined by challenging the mice with 6.0×10^9^ CFU *S. suis* strain SC19. Eight HIPs provide different levels of protection against challenge. Asterisks indicate that the HIPs are toxic and caused the death of the mice during boost immunization (see Fig.S5 for the details).

Challenge data showed that 7 of these HIPs, namely 5995, 6155, 2085, 575, 3685, 10025, and 10530 can provide partial protection (20%-40%) to immunized mice against challenge with the highly virulent *S. suis* SC19 strain (Fig. 5C). Significantly, HIP 8455 can provide complete protection against challenge compared to other HIPs. None of the remaining HIPs protected mice from challenge, despite being effective in inducing IgG antibodies. We found that HIPs 565, 2130, 5805, 6615, 10530, and 8455 were toxic and caused death in mice during boost immunization (Fig. S5). HIP 10025 has been tested in combination with 5 other *S. suis* proteins as a vaccine candidate in pigs, but no significant protection was observed after intranasal challenge with *S. suis* serotype 14 (36). The HIP 5995 (known as PrsA), which was shown to provide 50%-66% protection against challenge with *S. suis* type 2 or 9 strains in the previous study (35), induced 40% immune protection in our study. Overall, these results were consistent with previous studies, whereas the difference in efficacy of these HIPs observed in the current study may be due to differences in challenge dose and bacterial strain. Indeed, strain SC19, isolated from a diseased piglet during the 2005 human *S. suis* outbreak, is a hypervirulent strain that causes streptococcal toxic shock-like syndrome in humans, mice, and pigs (44, 45). Our results indicated that 8 HIPs are protective antigens of *S. suis* and can be used individually or in combination for the development of *S. suis* vaccines, particularly 8455. However, further work is needed to eliminate the toxicity of 8455 while maintaining its immunogenicity.

## Discussion

Identification of the conserved protective antigens across different serotypes is a game changer for the development of a broadly effective bacterial vaccine. Here, we developed a novel unbiased genome-wide approach, called BacScan, to identify the HIPs conserved across different serotypes by combining phage display, phage-immunoprecipitation, phage-PCR, and next-generation sequencing. A number of features make BacScan one of the most powerful approaches for identifying conserved immunogenic proteins for vaccine or diagnostic purposes. First, BacScan is an unbiased genome-wide screening approach. The genome-wide display library was constructed using T7 instead of filamentous phage, which is assembled in the periplasm of *E. coli* and only displays the peptides/proteins that are transported to the periplasm (12, 13). This is not the case for the lytic T7 phage that assembles in the cytoplasm (32). Most importantly, BacScan requires only one-round of panning instead of the multiple rounds of panning required by traditional phage selection (46, 47). This is important because the expression of certain exogenous proteins can limit the proliferation of the recombinant phages. As a result, these phages are lost after multiple rounds of amplification. In addition, all *S. suis* genes are split into equal length 600 bp fragments to minimize amplification bias during PCR amplification of the enriched phages (see Materials and Methods for the details).

Second, BacScan is not limited by the temporal expression of bacterial proteins. The expression of bacterial proteins is highly regulated, and not all immunogenic proteins are expressed when bacteria are cultured *in vitro* (48). Unlike some proteomics methods that use bacteria cultured *in vitro* to identify bacterial immunogenic proteins, BacScan used T7 phage to heterologously express and display all the core proteins of the pathogenic bacterium in *E. coli*. In addition, each protein was split into 200-aa fragments to avoid the possible expression issues with the full-length proteins, and therefore each protein has a higher chance of expressing at least one fragment. Therefore, all the bacterial proteins included in the T7 phage library were expected to be expressed.

Third, BacScan is a cost-effective method. Protein array is a high-throughput technology that allows for the identification of bacterial antigenic proteins (49, 50). However, the production of thousands of proteins to prepare protein arrays is very expensive, which is a limitation for some research laboratories. Reverse vaccinology reduces the number of bacterial candidate proteins, but still requires the expression of hundreds of proteins for their immunogenicity testing (9, 51). In contrast, BacScan does not require individual purification of bacterial proteins. Instead, the protein-encoding genes are cloned into T7 phage genome and expressed during phage propagation, which is much less expensive. In addition, the foreign bacterial genes in the genome of the enriched T7 phages are amplified by PCR, where different barcodes can be included for multiple serum samples used for antigen screening. The PCR results can be pooled for next-generation sequencing, which further reduces the cost.

As a proof of concept, we used BacScan to scan the *S. suis* core genome for the highly immunogenic proteins. All the 19 HIPs identified are highly conserved among different *S. suis* serotypes with 84-98% sequence identity. We found that 8 conserved HIPs can confer 20-100% protection to immunized mice against challenge with the highly virulent *S. suis* SC19 strain. Six of them, namely 6155, 2085, 575, 3685, 10530, and 8455, have not been previously tested as vaccine antigens. Significantly, HIP 8455 can provide complete protection against challenge, suggesting that it may be a good target for the development of a broadly protective subunit vaccine. In addition, we found that two conserved HIPs, 2085 and 6155, which have low homology with proteins from other bacteria, could be specific targets for the development of serological techniques for the diagnosis of *S. suis*.

Although BacScan is a powerful technology for identifying conserved immunogenic proteins, it has several limitations. First, the identification of protective antigens is highly dependent on the quality of the serum. If the serum cannot neutralize bacterial cells, it is impossible to screen for bacterial protective HIPs using BacScan. Therefore, it is strongly recommended that the neutralizing activity of the serum be determined prior to screening. Second, the bacterial proteins in the T7 display library were split into 200-amino acid long fragments, which may disrupt the structure of some proteins. This can be overcome by using multiple T7 libraries displaying different lengths of proteins and each library displaying proteins of the same length.

In summary, we have developed an unbiased and genome-wide approach, BacScan, to scan the bacterial core genome for the highly immunogenic proteins. This technology can cost-effectively identify conserved antigenic proteins across different serotypes, and we believe it will accelerate the development of a broadly protective subunit vaccine.

## Materials and Methods

### Ethics statement

Animal experiments were performed in accordance with animal welfare principles and approved by the Research Ethics Committee Huazhong Agricultural University, Hubei, China (HZAUMO-2023-0068). The studies of human sera were reviewed and approved by the Ethic Committee Tongji Medical College, Huazhong University of Science and Technology, Hubei, China (2023S080).

### Bacterial strains and phage

*S. suis* strain SC19 was cultured in tryptic soy broth or on tryptic soy agar containing 10% newborn bovine sera. *E. coli* strains DH5α (*hsdR17(rK− mK+) sup^2^*) and BL21 (DE3) were used for plasmid construction and protein expression, respectively. T7 phage was used to construct the display library as previously described (31). *E. coli* BLT5615 was used as host cells for the propagation of T7 display library.

### Serum samples

Six-week-old female BALB/c mice were obtained from the Laboratory Animal Center of Huazhong Agricultural University, Hubei, China, for the preparation of S. suis hyperimmune sera. The mice were randomly divided into two groups. The mice were infected intraperitoneally (n=5) with 4×10^7^ CFU *S. suis* SC19 strain three times on days 0, 14, and 21. Mice (n=5) receiving 200 μL PBS were used as negative controls. Blood samples were collected from the tail vein 7 days after the third infection. Clinical pig sera (n=20) and *S. suis* negative pig sera (n=20) were obtained from the Animal Disease Diagnostic Center of Huazhong Agricultural University, Hubei, China. Sixty serum samples were collected from farm workers with a history of close contact with pigs. Of these, 46 had a history of close contact with pigs for >5 years and 14 had ≤5 years of exposure. Twenty serum samples from children with no history of close contact with pigs were used as controls.

### Construction of T7 phage display library

The core genes of *S. suis* were identified as described previously (52) and amplified by PCR using gene-specific primers from *S. suis* strain SC19 genomic DNA. The genes over 600 bp were split into 600 bp fragments with 300 bp overlap between adjacent fragments. If the last fragment of a gene is less than 600 bp, the fragment was extended to 600 bp towards to the 5’ end of the gene. For the genes smaller than 600 bp, the fragment was extended to 600 bp towards to the next gene. The PCR products were digested with EcoRI/HindIII or BamHI/NotI depending on the presence of endonuclease recognition sites in the genes. Generated DNA fragments were then ligated into the T7 10-3b vector (Millipore) linearized with the same endonucleases to generate T7 gp10A-*S. suis* fragment fusion genes. The ligation products were packaged into T7 phage capsid *in vitro* using T7 packaging extracts and the generated recombinant T7 phages were propagated in *E. coli* BLT5615.

### Phage immunoprecipitation

Phage immunoprecipitation was performed as described previously (53). Briefly, 1.5 ml microcentrifuge tubes were blocked with 3% BSA at 4°C overnight. Mouse serum containing 2μg IgG and 2×10^9^ pfu phages was mixed in 1.5ml microcentrifuge tube. After 18h incubation at 4℃ with rotation, 40 μl of protein A/G magnetic beads (Thermo Fisher Scientific) were added and further incubated for 4h at 4℃. The mixture was centrifuged at 900×g for 1 minute at room temperature, and the tube was then placed onto a magnetic stand to remove the supernatant. The magnetic beads were resuspended with 600 μl IP wash buffer (50 mM Tris-HCl, pH 7.5, 150 mM NaCl, 0.1% NP-40) and transferred to a new 1.5ml microcentrifuge tube. After two washes with 1mL of IP wash buffer, the magnetic beads were resuspended with 40 μl deionized water, and the coprecipitated phages were lysed at 95℃ for 10 minutes. Mouse IgG concentration was determined using the bovine serum albumin (BSA) as a standard. All samples were performed in triplicate.

### Phage PCR and sequencing

The *S. suis* gene fragments were amplified from phage lysis using the primers F1 (5’-TTGT CTTCCTAAGACCGCTTGGCCTCCGACTTGGGGTTAACTAGTTACTCGAGTGCGG-3’) and R1 (5’-CCGAACGCAGCAAACTACGC-3’). The generated PCR products were used as templates for the second round PCR using primers F2 (5’-GAACGACATGGCTACGATCCGACTTTCGTATTCCAGTCAGGTGTGATGCTCGG-3’) and R2 (5’-TGTGAGCCAAGGAGTTGxxxxxxxxxxTTGTCTTCCTAAGACCGCTTGGCCT-3’, the “xxxxxxxxxx” represents 10 nt MGI regular index sequence to distinguish different samples). PCR products were purified by agarose gel electrophoresis and sequenced on a MGISEQ-2000 NextSeq platform (BGI) at the National Key Laboratory of Crop Genetic Improvement, Huazhong Agricultural University. Low-quality sequencing data were filtered out using fastp (ver 0.20.0), and the PCR amplification-induced adapter sequences were removed using cutadapt (ver 1.18) with the default settings (54). The resulted clean data were used to identify the highly immunogenic proteins as described previously (53).

### Bioinformatic analysis of the HIPs

The function of 793 core proteins of *S. suis* and 12 reported reference proteins were annotated in the Eggnog5 database (http://eggnog5.embl.de/#/app/home), and the members in each category were counted before and after screening. The enrichment of each category was determined by Fisher Exact Test. The virulence-associated genes (VAGs) of the *S. suis* in the 805 genes were identified by local blastp program using the VAG database described in the previous study (43). The enrichment of VAGs after screening was analyzed using Fisher Exact Test. Sequence similarity of HIPs with other proteins was determined using NCBI blastp online tool. The reference strains of six different serotypes *S. suis* were randomly selected to calculate the sequence conservation of HIPs. Nine *streptococcal* species commonly found in human clinics were selected to analyze the conservation of HIPs among genera. Extra-*Streptococcus* species were used to identify the homologies of *S. suis* HIPs in other bacteria. Blast results with coverage less than 80% coverage were ignored. The homology data were shown on a heatmap generated by the R package “pheatmap”, and the “Out Genus” part of the heatmap only included species homologous with at least 5 HIPs.

### Expression and purification of recombinant HIPs

The genes encoding highly immunogenic proteins (HIPs) were amplified from the genome of strain SC19 and cloned individually into linearized pET-28a or pET-32a vector. The generated plasmid was transformed into *E. coli* Bl21 (DE3), and the expression of recombinant protein was induced with 1mM isopropyl-β-D-thiogalactoside at 30°C for 4 hours. The *E. coli* cells were harvested by centrifugation at 8,000×g for 10 min at 4℃ and lysed by high-pressure cell disruptor at 4°C. Cell debris was removed by centrifugation at 34,000×g at 4 °C for 22 min, and the supernatant was filtered through 0.22 μm filters. The recombinant proteins were purified using a Ni-NTA column (Yeasen, Wuhan, China) and concentrated by ultrafiltration (Millipore). Protein concentration was determined by SDS-PAGE using BSA as a standard.

### Antibody dynamic analysis of HIPs

Eight-week-old female BALB/c mice were divided into 3 groups (n=5) and intranasally infected once with 2.7 ×10^7^, 8 ×10^7^, and 2.4 ×10^8^ CFU of *S. suis* SC19 strain, respectively. To determine the effect of age on antibody dynamics, four-week-old (n=5) and six-week-old (n=5) female BALB/c mice were intranasally infected with 8 ×10^7^ CFU of *S. suis* SC19. Sera were collected from the tail vein at different time points. Antigenic-specific IgG titer were determined by indirect ELISA as described below.

### Enzyme-linked immunosorbent assay (ELISA)

Antigenic-specific antibody titers were determined by indirect ELISA as described previously (55). Briefly, the ELISA plates were coated with 100 ng of purified protein diluted in 100μL sodium carbonate buffer (pH 9.6) at 4°C overnight, and then blocked with 3% BSA in 200 μL PBST (PBS containing 0.05% Tween-20) at 37 ℃ for 1 hour. After five washes with PBST, 100 μL of PBST-diluted mouse, pig, or human sera were added to each well. The plate was incubated at 37 ℃for 1 hour followed by five washes with 250μL PBST. The secondary antibodies (HRP-conjugated goat anti-mouse IgG, HRP-conjugated goat anti-pig IgG, HRP-conjugated goat anti-human IgG) were diluted 1:5000 in PBST and added to each well. After 50 minutes incubation at 37°C, the plates were washed five times with PBST. 100μL of TMB (3,3′,5,5′-tetramethylbenzidine) substrate solution was added to each well and incubated for 5 minutes at room temperature. The reactions were stopped with 2 M H_2_SO_4_, and the plates were detected with a microplate reader at an absorbance of 450nm.

### Mouse immunization and challenge

Six-week-old female BALB/c mice were randomized into 17 groups (n=5) and immunized intraperitoneally with either 30μg purified protein in 100μL PBS emulsified with an equal volume Montanide ISA-201 (SEPPIC, France) or 100μL PBS control emulsified with 100μL Montanide ISA-201. All the mice were boosted 14 days after the first immunization. Serum samples were collected from the tail vein two weeks after the last immunization. For challenge, fresh cultured *S. suis* SC19 was collected by centrifugation (8,000×g) for 5 min at 4°C. Cell pellets were resuspended with PBS. Each mouse was infected intraperitoneally with 6×10^9^ CFU of *S. suis* SC19. All mice were continuously recorded for morbidity and mortality for one week.

### Statistical Analysis

Statistical analysis was performed by Student t-test using Prism Graphpad software, and multi-group comparisons were evaluated by one-way analysis of variance (ANOVA). In all cases, p<0.05 was considered statistically significant.

## Author contributions

Conceptualization: PT, JD; Methodology: JD, QZ, JY, YZ, ZM, SP, HQ; Supervision: AZ, GW, PT; Writing, review & editing: PT, JD, and AZ.

## Data and materials availability

All data in the study are available upon request from the authors.

## REFERENCES

1. G. B. D. A. R. Collaborators, Global mortality associated with 33 bacterial pathogens in 2019: a systematic analysis for the Global Burden of Disease Study 2019. Lancet 400, 2221–2248 (2022).

2. C. Antimicrobial Resistance, Global burden of bacterial antimicrobial resistance in 2019: a systematic analysis. Lancet 399, 629–655 (2022).

3. I. Bekeredjian-Ding, Challenges for Clinical Development of Vaccines for Prevention of Hospital-Acquired Bacterial Infections. Front Immunol 11, 1755 (2020).

4. C. G. Baums et al., Streptococcus suis bacterin and subunit vaccine immunogenicities and protective efficacies against serotypes 2 and 9. Clin Vaccine Immunol 16, 200–208 (2009).

5. S. Quessy, J. D. Dubreuil, M. Caya, R. Letourneau, R. Higgins, Comparison of pig, rabbit and mouse IgG response to Streptococcus suis serotype 2 proteins and active immunization of mice against the infection. Can J Vet Res 58, 220–223 (1994).

6. M. Coppola, T. H. Ottenhoff, Genome wide approaches discover novel Mycobacterium tuberculosis antigens as correlates of infection, disease, immunity and targets for vaccination. Semin Immunol 39, 88–101 (2018).

7. F. A. Bidmos, S. Siris, C. A. Gladstone, P. R. Langford, Bacterial Vaccine Antigen Discovery in the Reverse Vaccinology 2.0 Era: Progress and Challenges. Front Immunol 9, 2315 (2018).

8. S. J. Goodswen, P. J. Kennedy, J. T. Ellis, A guide to current methodology and usage of reverse vaccinology towards in silico vaccine discovery. FEMS Microbiol Rev 47 (2023).

9. M. Pizza et al., Identification of vaccine candidates against serogroup B meningococcus by whole-genome sequencing. Science 287, 1816–1820 (2000).

10. S. Plotkin, History of vaccination. Proc Natl Acad Sci U S A 111, 12283–12287 (2014).

11. B. R. Naidu et al., An immunogenic epitope of Chlamydia pneumoniae from a random phage display peptide library is reactive with both monoclonal antibody and patients sera. Immunol Lett 62, 111–115 (1998).

12. J. Rakonjac, M. Russel, S. Khanum, S. J. Brooke, M. Rajic, Filamentous Phage: Structure and Biology. Adv Exp Med Biol 1053, 1–20 (2017).

13. D. Gagic, M. Ciric, W. X. Wen, F. Ng, J. Rakonjac, Exploring the Secretomes of Microbes and Microbial Communities Using Filamentous Phage Display. Front Microbiol 7, 429 (2016).

14. A. Zhang, C. Xie, H. Chen, M. Jin, Identification of immunogenic cell wall-associated proteins of Streptococcus suis serotype 2. Proteomics 8, 3506–3515 (2008).

15. O. Vytvytska et al., Identification of vaccine candidate antigens of Staphylococcus aureus by serological proteome analysis. Proteomics 2, 580–590 (2002).

16. O. Finco et al., Identification of new potential vaccine candidates against Chlamydia pneumoniae by multiple screenings. Vaccine 23, 1178–1188 (2005).

17. B. Perch, P. F. Kristjansen, K. Skadhauge, [Human-pathogenic Group R streptococci. 2 cases of meningitis and one case of fatal sepsis]. Ugeskr Laeger 130, 1130–1132 (1968).

18. G. Goyette-Desjardins, J. P. Auger, J. Xu, M. Segura, M. Gottschalk, Streptococcus suis, an important pig pathogen and emerging zoonotic agent-an update on the worldwide distribution based on serotyping and sequence typing. Emerg Microbes Infect 3, e45 (2014).

19. J. P. Arends, N. Hartwig, M. Rudolphy, H. C. Zanen, Carrier rate of Streptococcus suis capsular type 2 in palatine tonsils of slaughtered pigs. J Clin Microbiol 20, 945–947 (1984).

20. C. Marois, L. Le Devendec, M. Gottschalk, M. Kobisch, Detection and molecular typing of Streptococcus suis in tonsils from live pigs in France. Can J Vet Res 71, 14–22 (2007).

21. A. van Samkar, M. C. Brouwer, C. Schultsz, A. van der Ende, D. van de Beek, Streptococcus suis Meningitis: A Systematic Review and Meta-analysis. PLoS Negl Trop Dis 9, e0004191 (2015).

22. M. Gottschalk, J. Xu, C. Calzas, M. Segura, Streptococcus suis: a new emerging or an old neglected zoonotic pathogen? Future Microbiol 5, 371–391 (2010).

23. Y. T. Huang, L. J. Teng, S. W. Ho, P. R. Hsueh, Streptococcus suis infection. J Microbiol Immunol Infect 38, 306–313 (2005).

24. J. Tang et al., Streptococcal toxic shock syndrome caused by Streptococcus suis serotype 2. PLoS Med 3, e151 (2006).

25. X. Dong et al., The global emergence of a novel Streptococcus suis clade associated with human infections. EMBO Mol Med 13, e13810 (2021).

26. C. Tan, A. Zhang, H. Chen, R. Zhou, Recent Proceedings on Prevalence and Pathogenesis of Streptococcus suis. Curr Issues Mol Biol 32, 473–520 (2019).

27. L. Liu et al., Identification and experimental verification of protective antigens against Streptococcus suis serotype 2 based on genome sequence analysis. Curr Microbiol 58, 11–17 (2009).

28. J. Li et al., Evaluation of the immunogenicity and the protective efficacy of a novel identified immunogenic protein, SsPepO, of Streptococcus suis serotype 2. *Vaccine* **29**, 6514–6519 (2011).

29. K. Huang et al., Identification and characterisation a surface-associated arginine peptidase in Streptococcus suis serotype 2. Microbiol Res 170, 168–176 (2015).

30. W. Li, X. Hu, L. Liu, H. Chen, R. Zhou, Induction of protective immune response against Streptococcus suis serotype 2 infection by the surface antigen HP0245. FEMS Microbiol Lett 316, 115–122 (2011).

31. G. J. Xu et al., Viral immunology. Comprehensive serological profiling of human populations using a synthetic human virome. Science 348, aaa0698 (2015).

32. S. Danner, J. G. Belasco, T7 phage display: a novel genetic selection system for cloning RNA-binding proteins from cDNA libraries. Proc Natl Acad Sci U S A 98, 12954–12959 (2001).

33. J. Seele et al., The immunoglobulin M-degrading enzyme of Streptococcus suis, IdeSsuis, is a highly protective antigen against serotype 2. *Vaccine* **33**, 2207–2212 (2015).

34. Y. Li et al., Identification of a surface protein of Streptococcus suis and evaluation of its immunogenic and protective capacity in pigs. Infect Immun 74, 305–312 (2006).

35. X. Jiang et al., Peptidyl isomerase PrsA is surface-associated on Streptococcus suis and offers cross-protection against serotype 9 strain. FEMS Microbiol Lett 366 (2019).

36. C. Weisse et al., Immunogenicity and protective efficacy of a Streptococcus suis vaccine composed of six conserved immunogens. Vet Res 52, 112 (2021).

37. A. Naz et al., Identification of putative vaccine candidates against Helicobacter pylori exploiting exoproteome and secretome: a reverse vaccinology based approach. Infect Genet Evol 32, 280–291 (2015).

38. A. A. Grassmann et al., Discovery of Novel Leptospirosis Vaccine Candidates Using Reverse and Structural Vaccinology. Front Immunol 8, 463 (2017).

39. V. Masignani, M. Pizza, E. R. Moxon, The Development of a Vaccine Against Meningococcus B Using Reverse Vaccinology. Front Immunol 10, 751 (2019).

40. J. Li et al., Reverse vaccinology approach for the identifications of potential vaccine candidates against Salmonella. Int J Med Microbiol 311, 151508 (2021).

41. R. S. Naorem et al., Identification of Putative Vaccine and Drug Targets against the Methicillin-Resistant Staphylococcus aureus by Reverse Vaccinology and Subtractive Genomics Approaches. Molecules 27 (2022).

42. E. Ong, M. U. Wong, Y. He, Identification of New Features from Known Bacterial Protective Vaccine Antigens Enhances Rational Vaccine Design. Front Immunol 8, 1382 (2017).

43. A. A. Estrada et al., Proposed virulence-associated genes of Streptococcus suis isolates from the United States serve as predictors of pathogenicity. Porcine Health Manag 7, 22 (2021).

44. L. Lin et al., An NLRP3 inflammasome-triggered cytokine storm contributes to Streptococcal toxic shock-like syndrome (STSLS). PLoS Pathog 15, e1007795 (2019).

45. L. Teng et al., Draft Genome Sequence of Hypervirulent and Vaccine Candidate Streptococcus suis Strain SC19. Genome Announc 5 (2017).

46. J. Li et al., Construction and characterization of a highly reactive chicken-derived single-chain variable fragment (scFv) antibody against Staphylococcus aureus developed with the T7 phage display system. Int Immunopharmacol 35, 149–154 (2016).

47. A. M. Piggott, A. M. Kriegel, R. D. Willows, P. Karuso, Rapid isolation of novel FK506 binding proteins from multiple organisms using gDNA and cDNA T7 phage display. Bioorg Med Chem 17, 6841–6850 (2009).

48. H. Gu, H. Zhu, C. Lu, Use of in vivo-induced antigen technology (IVIAT) for the identification of Streptococcus suis serotype 2 in vivo-induced bacterial protein antigens. BMC Microbiol 9, 201 (2009).

49. S. J. Lee et al., Identification of a common immune signature in murine and human systemic Salmonellosis. Proc Natl Acad Sci U S A 109, 4998–5003 (2012).

50. A. Olaya-Abril, I. Jimenez-Munguia, L. Gomez-Gascon, I. Obando, M. J. Rodriguez-Ortega, A Pneumococcal Protein Array as a Platform to Discover Serodiagnostic Antigens Against Infection. Mol Cell Proteomics 14, 2591–2608 (2015).

51. T. M. Wizemann et al., Use of a whole genome approach to identify vaccine molecules affording protection against Streptococcus pneumoniae infection. Infection and Immunity 69, 1593–1598 (2001).

52. L. A. Weinert et al., Genomic signatures of human and animal disease in the zoonotic pathogen Streptococcus suis. Nat Commun 6, 6740 (2015).

53. D. Mohan et al., PhIP-Seq characterization of serum antibodies using oligonucleotide-encoded peptidomes. Nat Protoc 13, 1958–1978 (2018).

54. A. J. E. von Hoyningen-Huene et al., Bacterial succession along a sediment porewater gradient at Lake Neusiedl in Austria. Sci Data 6 (2019).

55. M. Li et al., Bacteriophage T4 Vaccine Platform for Next-Generation Influenza Vaccine Development. Front Immunol 12, 745625 (2021).

