## Supplementary for "BacScan: An Unbiased and Genome-Wide Approach to Identify Bacterial Highly Immunogenic Proteins"

**Supplementary Figure 1.**

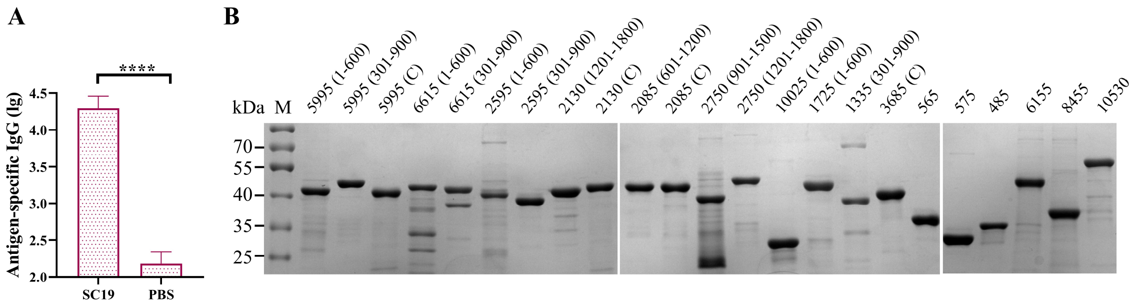


**Figure S1. Preparation of *S. suis*-specific sera and purified recombinant HIPs.**

**(A)** SPF mice were intraperitoneally infected with $4\times10^7$ CFU of *S. suis* strain SC19 three times on days 0, 14, and 21. Mice inoculated with the same volume of PBS were used as controls. Sera were collected 7 days after the third infection to determine *S. suis*-specific IgG by ELISA using inactivated SC19 cells as coating antigen. **(B)** SDS-PAGE analysis of purified recombinant proteins. A total of 24 S. suis gene fragments were enriched by BacScan and expressed in *E. coli*. Of these, 23 recombinant proteins can be purified and analyzed by SDS-PAGE.

**Supplementary Figure 2.**

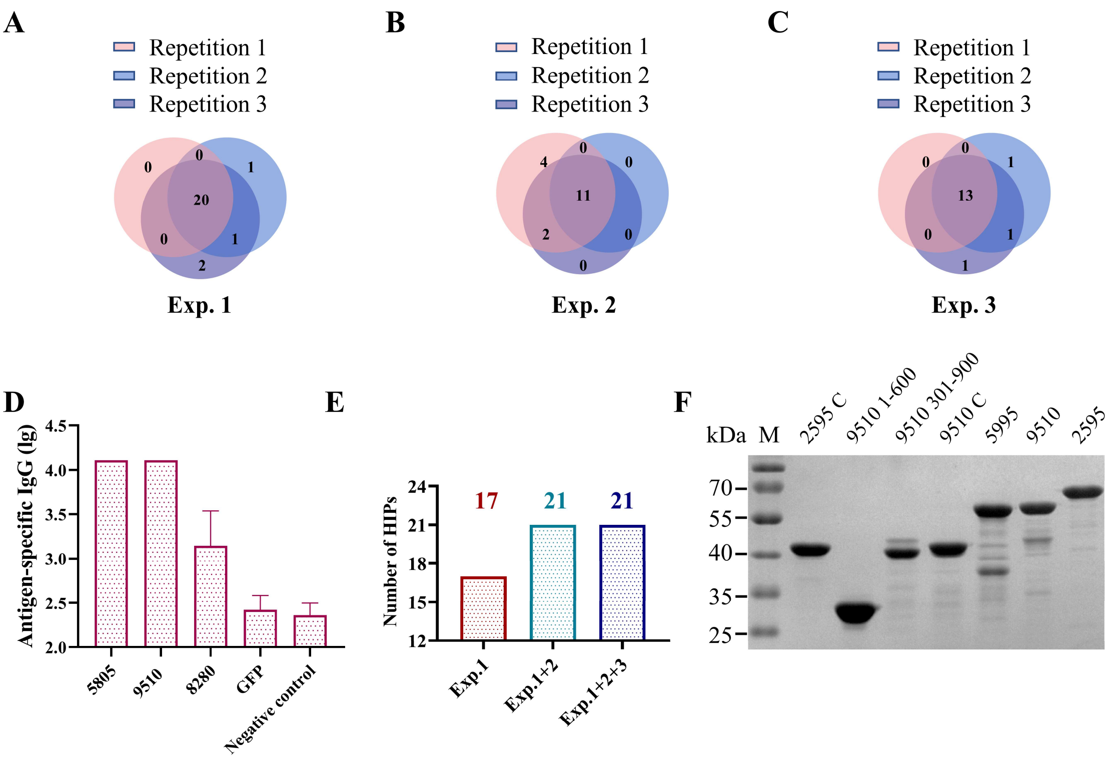


**Figure S2. Analysis of BacScan reproducibility**

Three independent BacScan screening experiments were performed on the same *S. suis* positive serum sample, and each of the independent experiments was performed in triplicate. A total of 24, 17, and 16 gene fragments were enriched in experiment #1 **(A)**, #2 **(B)**, and #3 **(C)**, respectively. The overlap of the enriched fragments in three parallel experiments in experiments #1 **(A)**, #2 **(B)**, and #3 **(C)** were shown. Compared to experiment #1, four novel HIPs were enriched in experiments # 2 and #3. Among them, 3 recombinant proteins were successfully purified and their binding activities to *S. suis*-specific sera were determined using ELISA **(D).** Recombinant GFP protein and *S. suis*-negative mouse sera were used as controls. **(E)** The number of HIPs identified for the same serum sample becomes increasingly saturated as the number of screens increases. A total of 17 HIPs were screened in experiment #1 and four novel HIPs were enriched in experiment #2, while no novel HIP was enriched in experiment #3. **(F)** Identification of immunodominant regions of HIPs using BacScan. The full-length HIP proteins and their fragments were purified and analyzed by SDS-PAGE.

**Supplementary Figure 3.**

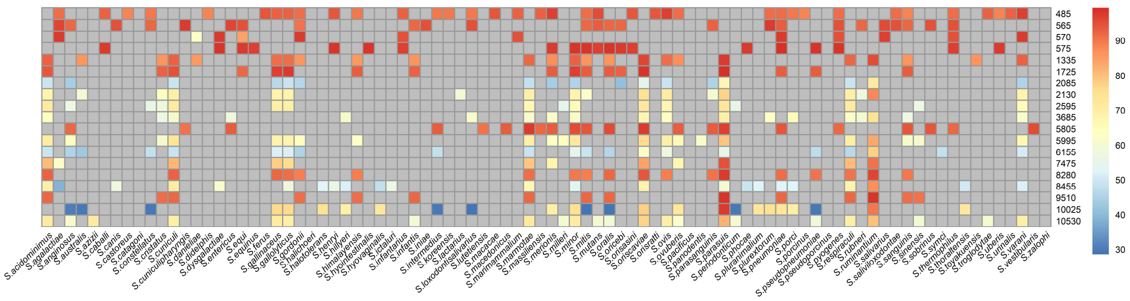


**Figure S3. Heatmap showing the conservation of 19 *S. suis* HIPs with the homology in Streptococcus.**

Sequence similarity analysis was performed using the NCBI blastp online tool. The conservation of 19 *S. suis* HIPs with the homology in 9 *streptococcal* species commonly found in human clinics was shown in Fig. 2D. The homology of 19 *S. suis* HIPs to proteins in the remaining *Streptococcus* species is shown here. Blast results with coverage less than 80% coverage were ignored. The heatmap was generated by the R package “pheatmap”. The color scale represents the identity (%) of each protein. All gray cells represent less than 80% coverage or no homologous proteins.

**Supplementary Figure 4.**

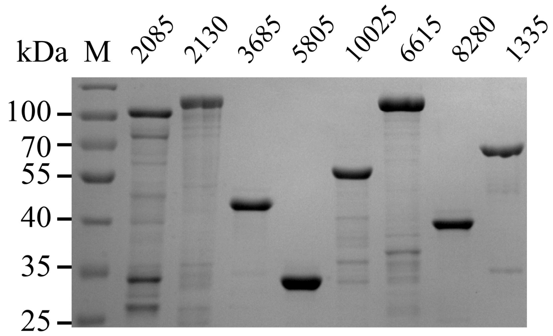


**Figure S4. SDS-PAGE analysis of the purified recombinant proteins.**

Enriched genes were expressed in *E. coli* Bl21 (DE3) and purified using HisTrap column as described in the Materials and Methods.

**Supplementary Figure 5.**

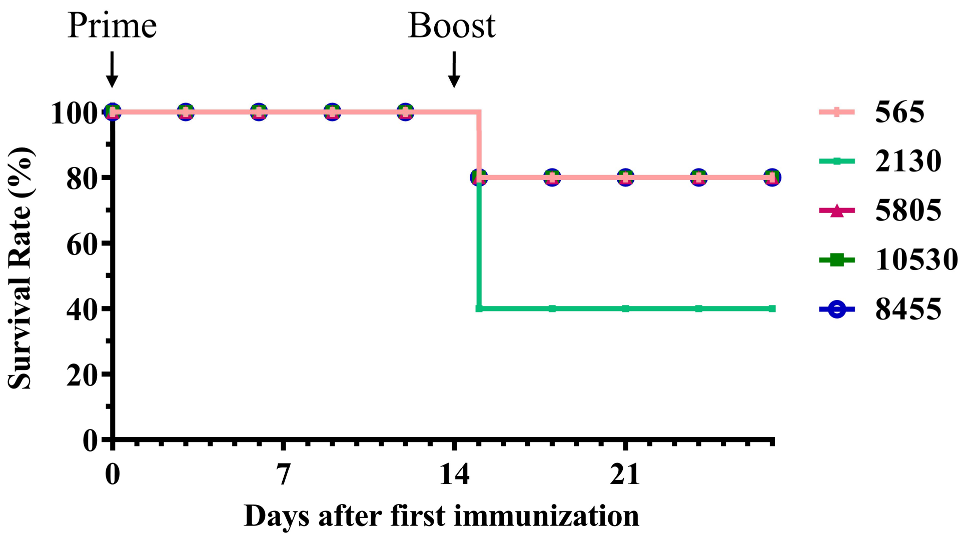


**Figure S5. Boost immunization of some HIPs caused death of mice**

The recombinant HIPs were emulsified with an equal volume of montanide ISA-201 and injected intraperitoneally into mice on days 0 and 14. Mice were monitored for mobility and mortality. Twenty percent of the mice died after boosted immunization with the 565, 5805, 10530 and 8455 proteins, while 60% of the mice died after boosted immunization with the 2130 protein.

**Table S1. The accession number of HIPs identified by BacScan in three independent experiments**

| Highly immunogenic proteins (HIPs) | Accession number | Identified in Exp # |
|---|---|---|
| 570 | ARL69119 | 1 |
| 575 | ARL69120 | 1 |
| 485 | ARL69102 | 1 |
| 6155 | ARL70113 | 1 |
| 8455 | ARL70545 | 1 |
| 1335 | ARL69242 | 1 |
| 2130 | ARL69386 | 1, 3 |
| 1725 | ARL69316 | 1, 3 |
| 2085 | ARL69378 | 1, 2 |
| 5805 | ARL70048 | 2, 3 |
| 8280 | ARL70512 | 2, 3 |
| 9510 | ARL70747 | 2, 3 |
| 7475 | ARL70357 | 2 |
| 10530 | ARL70936 | 1, 2, 3 |
| 565 | ARL69118 | 1, 2, 3 |
| 5995 | ARL70083 | 1, 2, 3 |
| 10025 | ARL70843 | 1, 2, 3 |
| 3685 | ARL69656 | 1, 2, 3 |
| 2595 | ARL69462 | 1, 2, 3 |
| 6615 | ARL70199 | 1, 2, 3 |
| 2750 | ARL69492 | 1, 2, 3 |
